# Bayesian Analysis for 3D combinatorial CRISPR screens

**DOI:** 10.1101/2022.06.02.494493

**Authors:** Lukas Madenach, Caroline Lohoff

## Abstract

Combinatorial CRISPR screens are a well-established tool for the investigation of genetic interactions in a high-throughput fashion. Currently, advancements from 2D combinatorial CRISPR screens towards 3D combinatorial screens are made, but at the same time an easy-to-use computational method for the analysis of 3D combinatorial screens is missing. Here we propose a Bayesian analysis method for 3D CRISPR screens based on a well-established 2D CRISPR screen analysis protocol. With our tool we hope to provide researchers with an out-of-the-box analysis solution, avoiding the need for time-consuming and resource-intensive development of custom analysis protocols.

## Introduction

Investigation techniques for the identification of genetic interactions in cancer with CRISPR screens has seen a lot of development over the recent years [1]. Currently advancements are made towards 3D genetic interaction screens, targeting three-way interactions, to gain new insights into complex genetic interaction networks [2]. However, a robust, go-to analysis method for 3D genetic interaction CRISPR screens is still lacking today. Here we propose the use of variational Bayesian inference techniques, which was already very sucessfully used for 2D genetic interaction CRISPR screens [3].

## Background

Genetic interactions are commonly defined as a combination of genetic alterations of at least two genes resulting in an effect greater than the additive effect of the individual contributing genetic alterations [4]. Genetic interactions have been studied through multiple techniques for example by investigating model systems such as yeast [5-7]. However, investigating genetic interactions in human cells on a larger scale required new methods, mainly because previous techniques for example those used in yeast are hard to scale to the sheer number of possible genetic interactions in human cells. An established, very powerful technique that enables high-throughput genetic interaction screening in human cells are combinatorial CRISPR screens, allowing researchers to find pairwise genetic interactions [8-10]. Pairwise genetic interaction analysis has led to a number of treatments for different cancers, such as the well-known treatment of BRCA1/2 mutated cancers, for example breast cancer, with PARP inhibitors [11]. Recently, in an effort to study more complex interactions, another step forward has been made towards 3D combinatorial CRISPR screens, allowing researchers to investigate the interaction among three genes in a high-throughput fashion [2].

Just as the techniques for conducting a combinatorial CRISPR screen have improved over the years, so have the analysis methods. One of the most advanced methods for analysing 2D combinatorial CRISPR screens, called GEMINI, uses a Bayesian approach to integrate gene effects and reagent effects as well as all samples and all replicates to infer the effects of interest [3]. While GEMINI is tailored for 2D combinatorial CRISPR screens and cannot be used for 3D combinatorial CRISPR screens directly, its general concept can be extended. Here we introduce Gemi3, a dedicated analysis method for 3D combinatorial CRISPR screens. We use a Bayesian approach to capture the effects of interest on the gene level as well as on the sgRNA level, while integrating available information from all samples and replicates. Additionally, Gemi3 is able to operate on a number of differently designed CRISPR screens, including asymmetrical designs.

## Methods

In an effort to make Gemi3 compatible with all 3D CRISPR screen pipelines, we use as input files a table containing the counts from the screen as well as a file with annotations for the guides (format requirements can be found in supplementary information).

### Bayesian model

We start by calculating the log-fold-change (LFC) of counts between two user-specified timepoints. To infer the sample-dependent and sample-independent effects from these LFC, we formulate them in a Bayesian framework and apply coordinate ascend variational inference (CAVI). We assume that the observed LFC can be broken down into four latent effects: the individual effect of each guide as well as the combinatorial effect of all three guides. Each of these effects can in turn be broken down into a sample-dependent effect and a sample-independent effect. We define the following notation:

- g, h and f – genes g, h and f being targeted simultaneously by one construct
- g_i_, h_j_ and f_k_ – guide i targeting gene g, guide j targeting gene h and guide k targeting gene f
- *l* – one sample
- 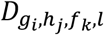 - observed LFC of guide triple (g_i_,h_j_,f_k_) in sample l

We assume the following model for the observed LFC:

- 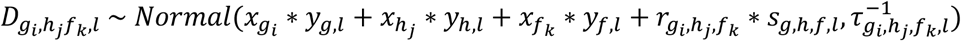

Together with prior distributions:

- 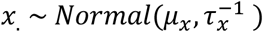
- 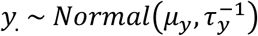
- 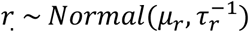
- 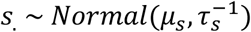
- *τ*. ∼ *Gamma*(*α, β*)

For the choices of prior distribution, we followed the recommendations made by Zamanighomi et al. [3]. The coordinate update equations needed for the CAVI algorithm were derived analytically for faster processing (supplementary information)[12]. After convergence of the CAVI algorithm, the inferred effects can be used to score the interactions.

### Scores

We implemented the following scoring system previously described elsewhere, although users can implement their own scoring system using the inferred effects [4].

The strong score is calculated independently for each sample and is defined as:

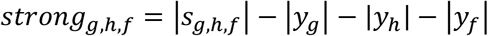

With this score we capture interactions that are „more than additive“. The larger the strong score, the larger the inferred interaction. A high strong score can be separated into two categories, lethality or recovery, depending on the sign of the respective *s*_*g,h,f*_ (Figure 1).

**Figure 1:**
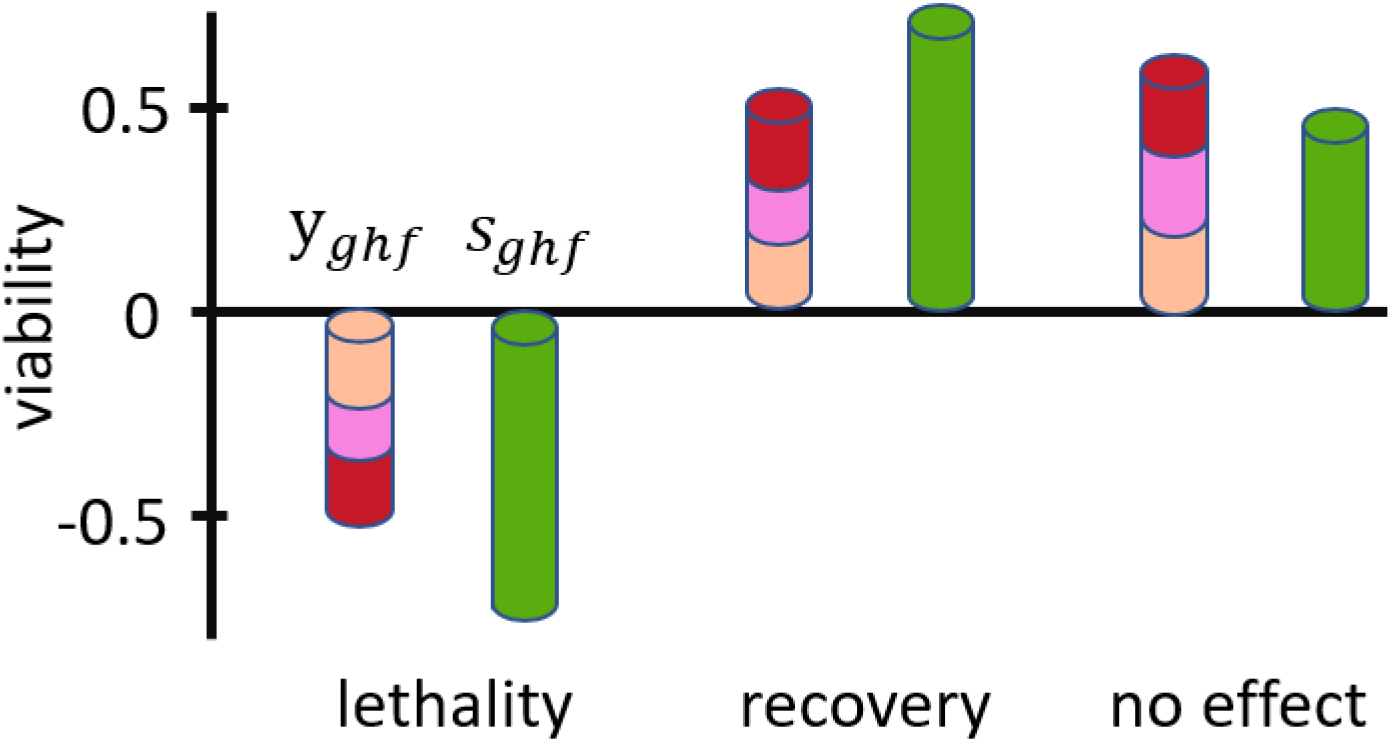
Types of genetic interaction captured by strong score. Direction of genetic interaction depends on the sign of s.

Further we implement sensitive recovery and sensitive lethality scores to capture more subtle interactions. The sensitive lethality score is calculated by the following formula where *pc*_*gene*_ means the y value of a provided positive control gene.

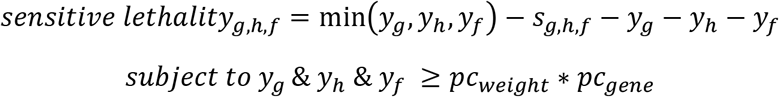

Analog to the sensitive lethality score, the sensitive recovery score is calculated.

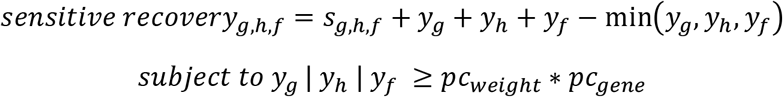

### Negative model

After all scores are calculated, a negative model is constructed for the calculation of p-values. For each score, all values associated with negative control genes are collected (Figure 2) and used to fit a gaussian mixture model (GMM).

**Figure 2:**
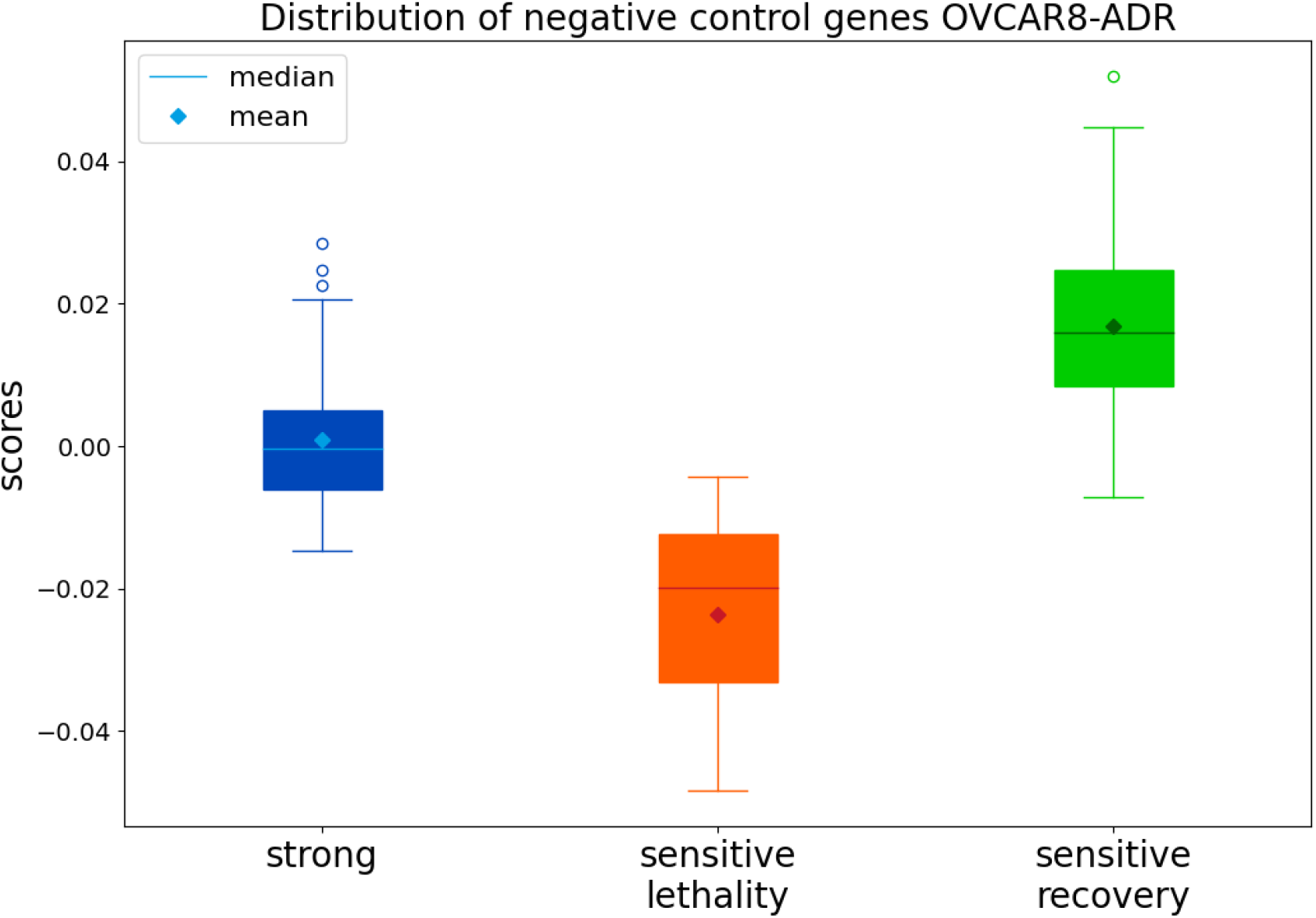
Distribution of scores associated with negative control genes for different score types.

**Figure 3:**
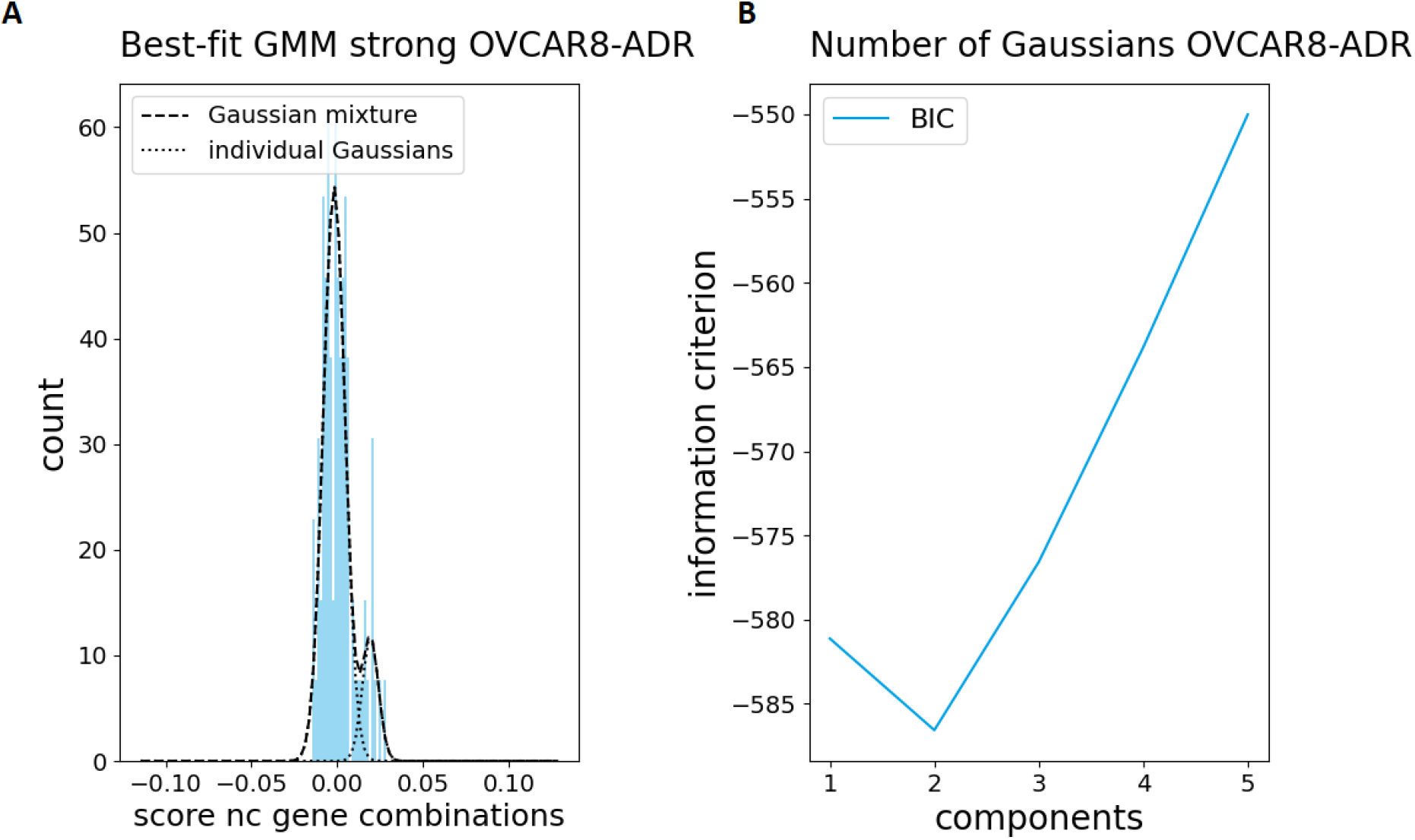
(A) GMM fitted to strong scores associated with negative control gene. (B) BIC used to select GMM. The smaller the BIC the better the fit of the model while simultaneously avoiding overfitting.

Up to five components are allowed for the GMM. The best GMM is chosen by Bayesian information criterion (BIC) [13], which controls for the number of used components in a GMM. Once fitted, the GMM is used as the null distribution to calculate p-values for the remaining scores. To account for multiple testing, p-values are corrected by the Benjamini-Hochberg method [14].

## Results

To demonstrate the utility of Gemi3 we used data from Zhou et al., to the best of our knowledge the only published 3D combinatorial CRISPR screen to date [2]. However, it should be mentioned, that the screen conducted by Zhou et al. was not set up with Gemi3 in mind and therefore lacks critical components such as a positive control gene and dedicated negative control genes in sufficient quantity. This heavily influences the calculated p-values. The 3D CRISPR screen was conducted in ovarian cancer cells (OVCAR8-ADR) with a total of 32768 different gRNA combinations. After obtaining the raw count data as described in Zhou et al. and annotation, the LFC was calculated and plotted. As expected, the LFC centers around zero (Figure 4).

**Figure 4:**
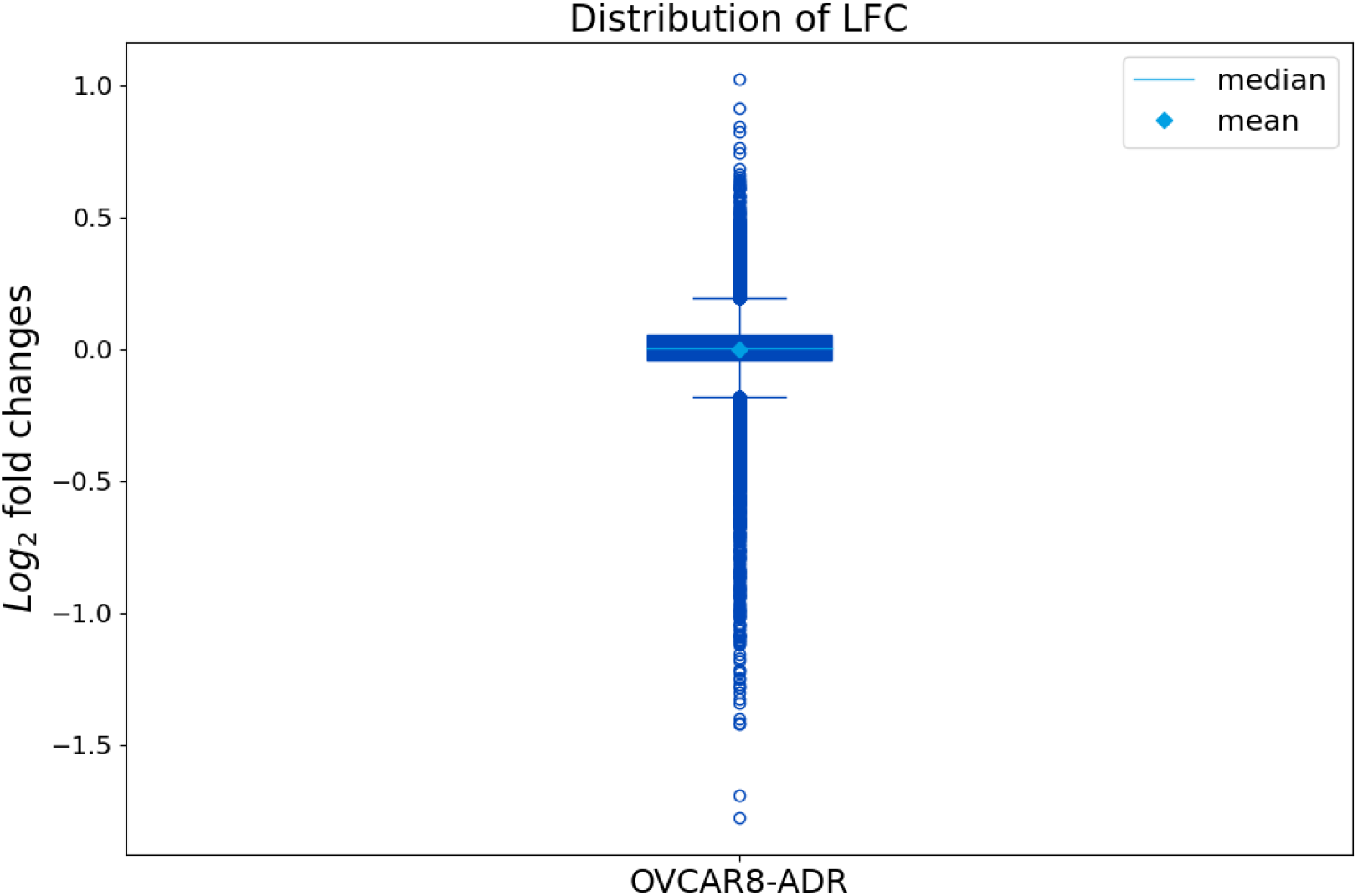
Distribution of all LFC values from Zhou et al.

After obtaining the LFC values the scores were calculated. The strong scores show the expected behavior, with the majority centered around zero and only a few combinations showing strong interaction (Figure 5).

**Figure 5:**
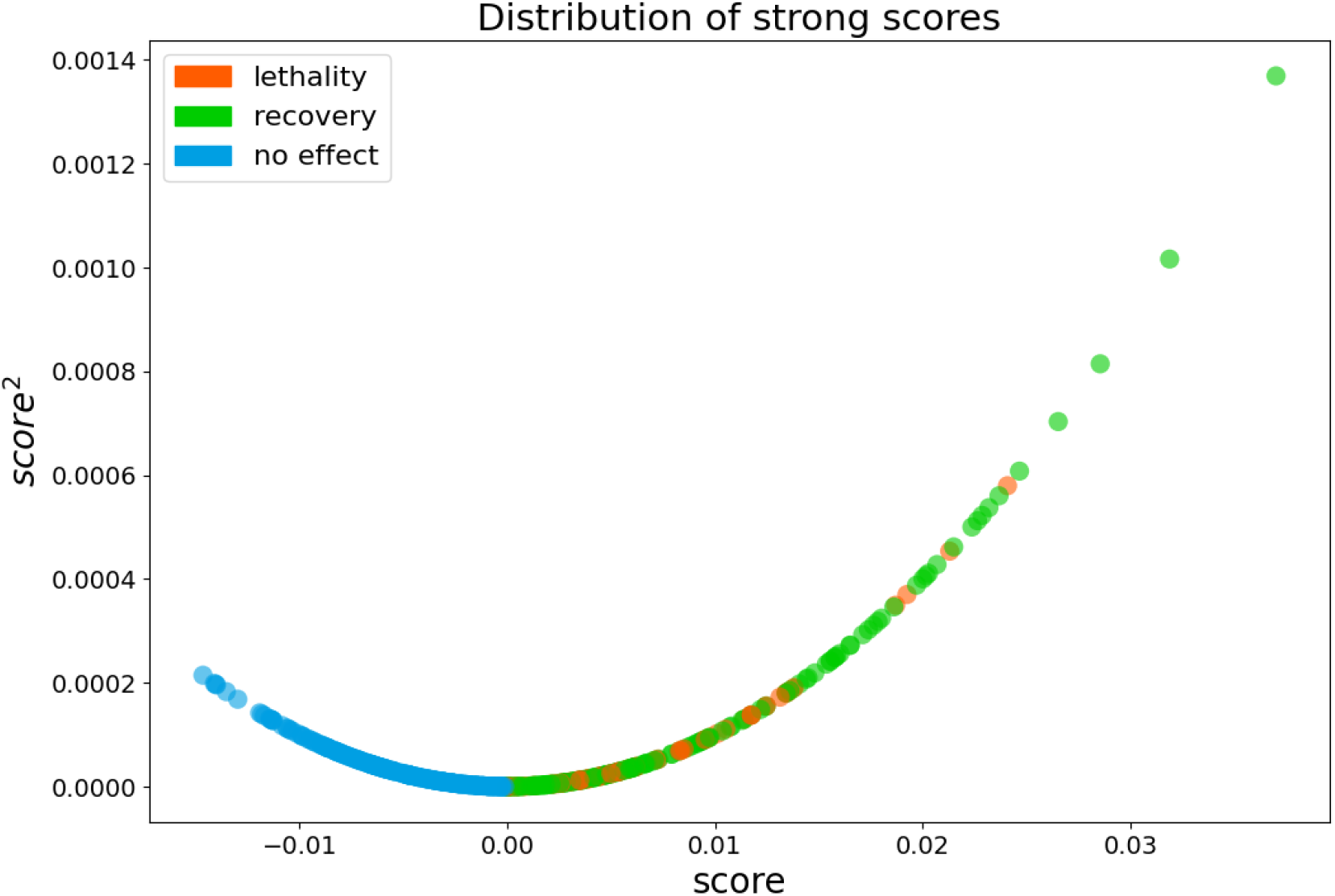
Distribution of strong score. Colors represent the direction of the underlying combinatorial effect.

To calculate p-values, scores associated with negative control genes were collected (Figure 2). For the strong scores the negative control genes center around zero as was expected. But for the sensitive lethality and sensitive recovery scores the negative control genes deviate from zero. This is most likely due to the fact that the screen was not set up with Gemi3 in mind and therefore lacks suitable negative control genes in sufficient quantity. A GMM was fitted on the distribution of scores associated with negative control genes to be used as null distribution for the calculation of p-values and FDR values (Figure 3). The strong scores together with their p-values are shown in a volcano plot as well as written to an output file (Figure 6).

**Figure 6:**
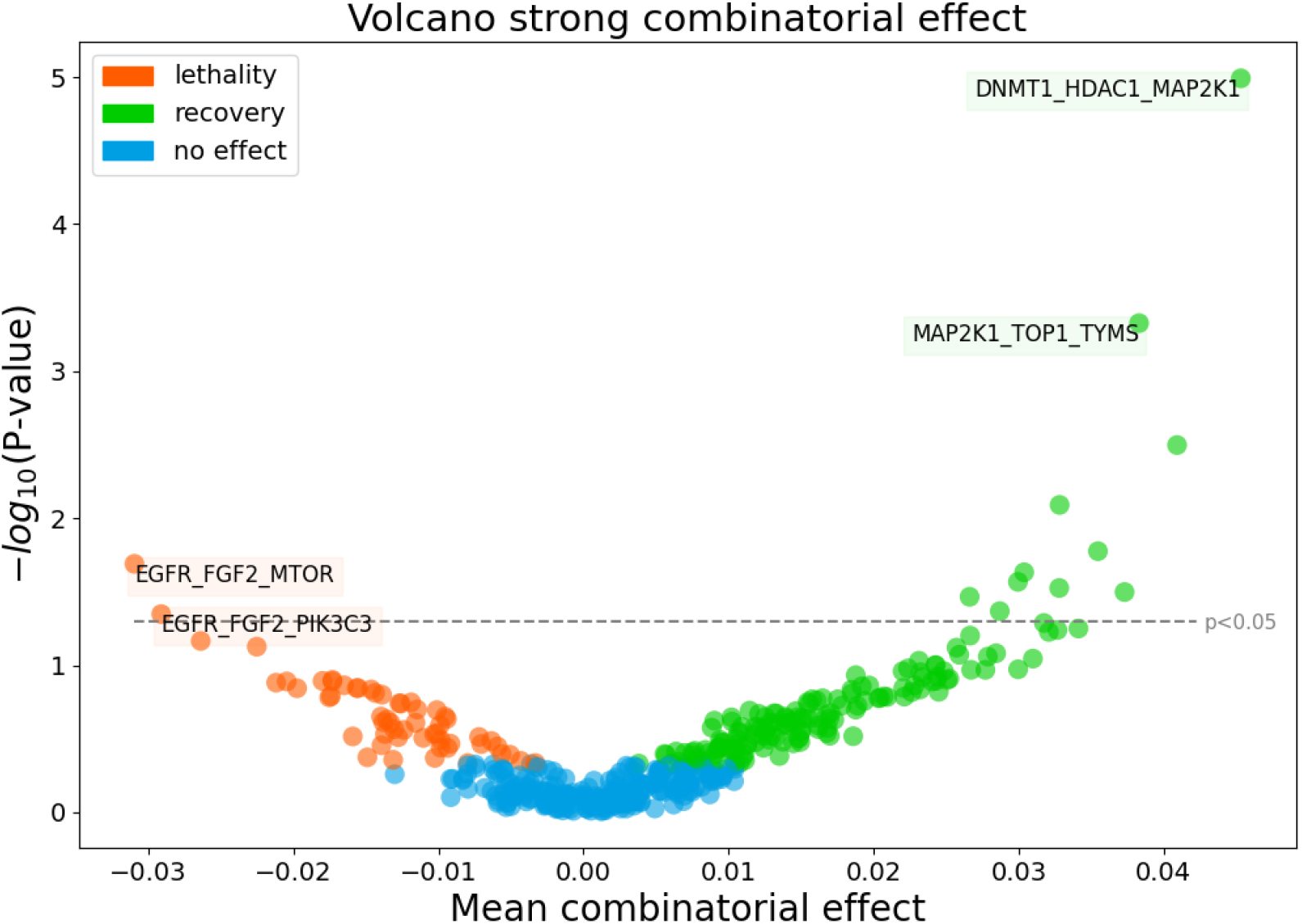
Results from the analysis with Gemi3. Combinatorial strong effects as well as statistical significance based on the GMM fitted to strong scores associated with negative control sgRNA is shown.

## Conclusion

Gemi3 is an analysis package dedicated to 3D combinatorial CRISPR screens, allowing for seamless use with existing CRISPR screen pipelines. It combines the user-friendliness and interoperability of Python with the high performance and scalability for HPC clusters of Rust. Additionally, to providing an out-of-the-box analysis method, Gemi3 also allows for highly customized analysis methods. While being usable for a variety of different 3D CRISPR screen concepts including asymmetrical designed CRISPR screens, one has to make sure that a sufficient number of control genes are included in the screen.

## Acknowledgements

We want to thank Dr. Natalie Jäger for helpful discussions and critical reading of the manuscript and Prof. Stefan M. Pfister for providing the fruitful environment for this project to take place.

## Availability

The described analysis method is implemented in the python package Gemi3 for user friendliness and seamless integration into existing processing pipelines while still offering high performance and scalability on HPC clusters due to the use of Rust for implementing the computation intensive CAVI algorithm. Code is available at https://github.com/Luk13Mad/Gemi3.

