## Supplementary Methods for "Bayesian Analysis for 3D combinatorial CRISPR screens"

### Gemi3 – Supplementary Information

#### Input files

Gemi3 requires two files as input. One file containing the raw counts from the screen (Figure 1). The file must be tab-separated and have a header line. As index (row names) the sgRNA sequences concatenated by a semicolon are used. The label for each row is the sample label. In the sample labels there must not be a „_“ character.

```
	OVCAR8-ADR
AAGCGAGT;AAGCGAGT;AAGCGAGT	0.5763439560976322
AAGCGAGT;AAGCGAGT;CTCTAGGT	0.3276618461252343
AAGCGAGT;AAGCGAGT;ACTACTCG	0.0543357385440228
AAGCGAGT;AAGCGAGT;ACGTGAGA	0.3465591328367694
AAGCGAGT;AAGCGAGT;AGTCCACA	0.3643811403582855
AAGCGAGT;AAGCGAGT;AGTTACAG	0.1399440267880121
AAGCGAGT;AAGCGAGT;ATAGGTGG	0.1628099397002707
AAGCGAGT;AAGCGAGT;ATCACGCA	0.147730354009433
AAGCGAGT;AAGCGAGT;ATTCAGCC	0.2458684235124548
AAGCGAGT;AAGCGAGT;ATTCGGTT	0.2044294820074244
AAGCGAGT;AAGCGAGT;CCTTCTCT	0.4803154660049822
AAGCGAGT;AAGCGAGT;CGAACTAG	0.3978165672177802
```

*Figure 1: First 13 lines of input file containing LFC data.*

The annotation file must also be tab-separated and have a header line (Figure 2). As index the same sgRNA sequences concatenated by a semicolon  as in the count file are used. Additionally the columns „gene_1“ , „gene_2“, „gene_3“, „guide_1“, „guide_2“ and „guide_3“ must be present, listing the sequence of the sgRNA and the associated gene names which are targeted by the respective guide.

```
rowname	gene_1	gene_2	gene_3	guide_1	guide_2	guide_3
AAGCGAGT;AAGCGAGT;AAGCGAGT	dummyguide1	dummyguide1	dummyguide1	AAGCGAGT	AAGCGAGT	AAGCGAGT
AAGCGAGT;AAGCGAGT;CTCTAGGT	dummyguide1	dummyguide1	dummyguide2	AAGCGAGT	AAGCGAGT	CTCTAGGT
AAGCGAGT;AAGCGAGT;ACTACTCG	dummyguide1	dummyguide1	CDK4	AAGCGAGT	AAGCGAGT	ACTACTCG
AAGCGAGT;AAGCGAGT;ACGTGAGA	dummyguide1	dummyguide1	CDK4	AAGCGAGT	AAGCGAGT	ACGTGAGA
AAGCGAGT;AAGCGAGT;AGTCCACA	dummyguide1	dummyguide1	DNMT1	AAGCGAGT	AAGCGAGT	AGTCCACA
AAGCGAGT;AAGCGAGT;AGTTACAG	dummyguide1	dummyguide1	DNMT1	AAGCGAGT	AAGCGAGT	AGTTACAG
AAGCGAGT;AAGCGAGT;ATAGGTGG	dummyguide1	dummyguide1	EGFR	AAGCGAGT	AAGCGAGT	ATAGGTGG
AAGCGAGT;AAGCGAGT;ATCACGCA	dummyguide1	dummyguide1	EGFR	AAGCGAGT	AAGCGAGT	ATCACGCA
AAGCGAGT;AAGCGAGT;ATTCAGCC	dummyguide1	dummyguide1	ERBB2	AAGCGAGT	AAGCGAGT	ATTCAGCC
AAGCGAGT;AAGCGAGT;ATTCGGTT	dummyguide1	dummyguide1	ERBB2	AAGCGAGT	AAGCGAGT	ATTCGGTT
AAGCGAGT;AAGCGAGT;CCTTCTCT	dummyguide1	dummyguide1	HDAC1	AAGCGAGT	AAGCGAGT	CCTTCTCT
AAGCGAGT;AAGCGAGT;CGAACTAG	dummyguide1	dummyguide1	HDAC1	AAGCGAGT	AAGCGAGT	CGAACTAG
```

*Figure 2: First 13 lines of annotation file.*

### CAVI - Analytically derived coordinate updates for posterior distributions

- g, h and f – genes g ,h and f being targeted simultaneously by one construct
- $g_i$, $h_j$ and $f_k$ – guide i targeting gene g, guide j targeting gene h and guide k targeting gene f
- l – one sample
- $D_{g_i,h_j,f_k,l}$ - observed LFC of guide triple ($g_i$,$h_j$,$f_k$) in sample l

Assuming the model:

$$D_{g_i,h_j f_k,l} \sim Normal(x_{g_i} * y_{g,l} + x_{h_j} * y_{h,l} + x_{f_k} * y_{f,l} + r_{g_i,h_j,f_k} * s_{g,h,f,l}, \tau^{-1}_{g_i,h_j,f_k,l})$$

Together with prior distributions:

$x_{.} \sim Normal(\mu_x, \tau_x^{-1})$

$y_{.} \sim Normal(\mu_y, \tau_y^{-1})$

$r_{.} \sim Normal(\mu_r, \tau_r^{-1})$

$s_{.} \sim Normal(\mu_s, \tau_s^{-1})$

$\tau_{.} \sim Gamma(\alpha, \beta)$

Where $\tau$ is the precision defined as $\tau = \frac{1}{\sigma^2}$. We approximate the posterior of the latent variables $(x, y, r, s \; and \; \tau)$ with the mean-field variational family, which assumes independence among the variables and factorizes over the variables.

$$q(x,y,r,s,\tau) = \prod_{g,i} q(x_{g_i}) * \prod_{g,l} q(y_{g,l}) * \prod_{g,i,h,j,f,k} q\left(r_{g_i,h_j,f_k}\right) * \prod_{g,h,f,l} q(s_{g,h,f,l}) * \prod_{g,i,h,j,f,k,l} q\left(\tau_{g_i,h_j,f_k,l}\right)$$

Note we don't make further assumptions about the form of the factors of $q(x,y,r,s,\tau)$. We find the optimal form for $q(x_{g_i})$ by evaluating the expression $E_{-x_{g_i}}[\ln f(D,x,y,r,s,\tau)]$ were $E_{-x_{g_i}}$ means the expectation value taken over all variables except $x_{g_i}$ . For a more detailed explanation see „Pattern recognition and machine learning“ – C. Bishop chapter 10 and following [1].

By evaluating the above expression for each variable we find the following closed expressions:

$$\boldsymbol{q(x_{g_i}) \sim Normal\left(\mu_{x_{g_i}}, \tau^{-1}_{x_{g_i}}\right)}$$

$$\mu_{x_{g_i}} = \frac{\tau_x * \mu_x + \sum_{h,j,f,k,l} E\left(\tau_{g_i,h_j,f_k,l}\right) * E(y_{g,l}) * \left[D_{g_i,h_j f_k,l} - E\left(x_{h_j}\right) * E(y_{h,l}) - E(x_{f_k}) * E(y_{f,l}) - E\left(r_{g_i,h_j,f_k}\right) * E(s_{g,h,f,l})\right]}{\tau_x + \sum_{h,j,f,k,l} E\left(\tau_{g_i,h_j,f_k,l}\right) * E(y^2_{g,l})}$$

$$\tau_{x_{g_i}} = \tau_x + \sum_{h,j,f,k,l} E\left(\tau_{g_i,h_j,f_k,l}\right) * E(y^2_{g,l})$$

$$\boldsymbol{q(y_{g,l}) \sim Normal\left(\mu_{y_{g,l}}, \tau_{y_{g,l}}^{-1}\right)}$$

$$\mu_{y_{g,l}} = \frac{\tau_y * \mu_y + \sum_{i,h,j,f,k} E\left(\tau_{g_i,h_j,f_k,l}\right) * E(x_{g_i}) * \left[D_{g_i,h_j f_k,l} - E\left(x_{h_j}\right) * E(y_{h,l}) - E(x_{f_k}) * E(y_{f,l}) - E\left(r_{g_i,h_j,f_k}\right) * E(s_{g,h,f,l})\right]}{\tau_y + \sum_{h,j,f,k,l} E\left(\tau_{g_i,h_j,f_k,l}\right) * E(x_{g_i}^2)}$$

$$\tau_{y_{g,l}} = \tau_y + \sum_{i,h,j,f,k} E\left(\tau_{g_i,h_j,f_k,l}\right) * E(x_{g_i}^2)$$

$$\boldsymbol{q\left(r_{g_i,h_j,f_k}\right) \sim Normal\left(\mu_{r_{g_i,h_j,f_k}}, \tau_{r_{g_i,h_j,f_k}}^{-1}\right)}$$

$$\mu_{r_{g_i,h_j,f_k}} = \frac{\tau_r * \mu_r + \sum_l E\left(\tau_{g_i,h_j,f_k,l}\right) * E(s_{g,h,f,l}) * \left[D_{g_i,h_j f_k,l} - E\left(x_{h_j}\right) * E(y_{h,l}) - E(x_{f_k}) * E(y_{f,l}) - E(x_{g_i}) * E(y_{g,l})\right]}{\tau_r + \sum_l E\left(\tau_{g_i,h_j,f_k,l}\right) * E(s_{g,h,f,l}^2)}$$

$$\tau_{r_{g_i,h_j,f_k}} = \tau_r + \sum_l E\left(\tau_{g_i,h_j,f_k,l}\right) * E(s_{g,h,f,l}^2)$$

$$\boldsymbol{q(s_{g,h,f,l}) \sim Normal\left(\mu_{s_{g,h,f,l}}, \tau_{s_{g,h,f,l}}^{-1}\right)}$$

$$\mu_{s_{g,h,f,l}} = \frac{\tau_s * \mu_s + \sum_{i,j,k} E\left(\tau_{g_i,h_j,f_k,l}\right) * E\left(r_{g_i,h_j f_k}\right) * \left[D_{g_i,h_j f_k,l} - E\left(x_{h_j}\right) * E(y_{h,l}) - E(x_{f_k}) * E(y_{f,l}) - E(x_{g_i}) * E(y_{g,l})\right]}{\tau_s + \sum_{i,j,k} E\left(\tau_{g_i,h_j,f_k,l}\right) * E\left(r_{g_i,h_j f_k}^2\right)}$$

$$\tau_{s_{g,h,f,l}} = \tau_s + \sum_{i,j,k} E\left(\tau_{g_i,h_j,f_k,l}\right) * E\left(r_{g_i,h_j f_k}^2\right)$$

$$\boldsymbol{q\left(\tau_{g_i,h_j,f_k,l}\right) \sim Gamma(\alpha_{\tau_{g_i,h_j,f_k,l}}, \beta_{\tau_{g_i,h_j,f_k,l}})}$$

$$\alpha_{\tau_{g_i,h_j,f_k,l}} = \alpha_{g_i,h_j,f_k,l} + \frac{1}{2}$$

$$\beta_{\tau_{g_i,h_j,f_k,l}} = \beta_{g_i,h_j,f_k,l} + \frac{1}{2} D_{g_i,h_j,f_k,l}^2 - D_{g_i,h_j,f_k,l} E(x_{g_i}) E(y_{g,l}) - D_{g_i,h_j,f_k,l} E\left(x_{h_j}\right) E(y_{h,l}) - D_{g_i,h_j,f_k,l} E(x_{f_k}) E(y_{f,l}) - D_{g_i,h_j,f_k,l} E\left(r_{g_i,h_j,f_k}\right) E(s_{g,h,f,l}) + E(x_{g_i}^2) E(y_{g,l}^2) + E(x_{g_i}) E(y_{g,l}) E\left(x_{h_j}\right) E(y_{h,l}) + E(x_{g_i}) E(y_{g,l}) E(x_{f_k}) E(y_{f,l}) + E(x_{g_i}) E(y_{g,l}) E\left(r_{g_i,h_j,f_k}\right) E(s_{g,h,f,l}) + \frac{1}{2} E\left(x_{h_j}^2\right) E(y_{h,l}^2) + E\left(x_{h_j}\right) E(y_{h,l}) E(x_{f_k}) E(y_{f,l}) + E\left(x_{h_j}\right) E(y_{h,l}) E\left(r_{g_i,h_j,f_k}\right) E(s_{g,h,f,l}) + E(x_{g_i}) E(y_{g,l}) E\left(r_{g_i,h_j,f_k}\right) E(s_{g,h,f,l}) + \frac{1}{2} E\left(r_{g_i,h_j,f_k}^2\right) E(s_{g,h,f,l}^2)$$

$$E\left(\tau_{g_i,h_j,f_k,l}\right) = \frac{\alpha_{\tau_{g_i,h_j,f_k,l}}}{\beta_{\tau_{g_i,h_j,f_k,l}}}$$

These closed expression are used to update one coordinate at a time while holding the other coordinates constant.
